# Laboratory mice with a wild microbiota generate strong allergic immune responses

**DOI:** 10.1101/2021.03.28.437143

**Authors:** Junjie Ma, Cajsa H. Classon, Julian M. Stark, Muzhen Li, Huey-Jy Huang, Susanne Vrtala, Stephan P. Rosshart, Susanne Nylén, Jonathan M. Coquet

## Abstract

Allergic disorders are caused by a combination of hereditary and environmental factors. The hygiene hypothesis postulates that early life microbial exposures impede the development of subsequent allergic disease. However, unambiguous evidence that microbes reduce the development of allergic disorders is still lacking. Recently developed ‘wildling’ mice contain a rich and diverse commensal and encounter a repertoire of microbes typical of the wild, with pathogenic potential. Here, we probed the hygiene hypothesis by comparing the development of allergic inflammation in wildlings to that of genetically identical mice lacking diverse microbial exposure. We find that wildlings develop stronger allergic inflammation in response to house dust mites with allergic T cell responses driven not only by cognate peptide antigens, but also by innate cytokines. In all, the results suggest that high microbial content and diversity potentiates, rather than restricts, allergic immune responses.

**One sentence summary:** Strong allergic inflammation in the face of rich and diverse microbial exposures

## Main text

The incidence of allergies has risen at an alarming rate over the past century (*1*). In some developed countries, approximately 30% of children are affected by rhinitis, atopic dermatitis or asthma by the age of 5 years (*2*). Genetic factors play an important role in susceptibility to allergic disease since concordance between monozygotic twins is approximately 50%. Genome wide association studies have pinpointed several loci involved in the development of allergy (*3*), including HLA-D alleles, which encode for major histocompatibility class II molecules.

However, the striking rise in allergic disorders during the 20^th^ century cannot solely be explained by genetic factors. In 1989, Strachan observed that children with several older siblings developed hay fever at considerably reduced rates to children that grew up with few or no siblings (*4*). He proposed that this was due to microbial transfer from older to younger siblings, and these findings were corroborated by similar studies of family size and sibship over the subsequent decade (*5*). Colloquially, Strachan’s findings came to be known as the *hygiene hypothesis* and were extrapolated to include other atopic disorders such as asthma and atopic dermatitis.

In support of Strachan’s observations, children that grew up on farms were found to have approximately half the incidence of asthma as those that grew up in urban settings (*6–8*). Sampling of the farm environment identified increased endotoxin exposure in animal barns, and the consumption of raw milk as factors that may reduce the incidence of asthma in children (*9–11*). Endotoxin exposure was subsequently shown to desensitize the airway epithelium to activation by pervasive allergens and reduce airway inflammation associated with asthma (*12*). Though the hygiene hypothesis is one plausible explanation for the rise in allergic disease in the 20^th^ century, several studies measuring environmental and household microbial exposures have failed to show a direct inverse correlation between exposure and allergy (*13–17*). There is a continued need for well-controlled investigations of the impact that the microbial environment exerts on allergic disease.

Preclinical studies of allergy have relied heavily on the use of mice that are housed under specific-pathogen-free (SPF) conditions. These have conclusively demonstrated that allergic immune responses are mediated by adaptive type 2 lymphocytes termed T helper 2 (Th2) cells and an analogous population of innate lymphoid cells (ILC2). A major mechanism by which microbial exposures are proposed to limit allergy is by subverting the immune response away from Th2 cell-mediated responses.

While laboratory mice have well defined genes and offer the potential for genetic manipulation, mice housed under SPF conditions do not come into contact with the breadth of pathogens present in the natural world and nor are they colonized by a rich and diverse, naturally co-evolved microbiome typically found in the natural world (*18, 19*).

Several recent studies show that mice housed under SPF conditions fail to faithfully replicate the immune responses of free-living humans (*19–22*). In studies of wild mice, pet store mice or SPF mice made wild by fecal material transfer or co-housing, researchers noted that these animals mounted more effective responses to influenza, bacteria, tumors and parasites than conventional laboratory SPF mice and better reflected the human situation (*19, 20*). Thus, although the microbial milieu is a central tenet of the hygiene hypothesis, the exclusive use of microbially-restrictive conventional laboratory SPF mice cannot adequately model the role that environmental factors may play in allergic disease. In the context of the hygiene hypothesis, the restricted commensal and pathogenic repertoire of microbes found in SPF mice might rather distort the development of the immune system, leading to false assumptions of how the human immune system functions.

Since direct comparisons between conventional laboratory mice and wild mice are complicated by genetic disparities, Rosshart and colleagues recently described a system for the transfer of common C57BL/6 mouse embryos into wild mouse surrogate mothers, generating so-called ‘wildling’ mice (*18*). Wildling mice were found to acquire a fulminant gut, skin and vaginal microbiota, a representative breadth of naturally occurring pathogens and many of the broader immune features of wild mice. Thus, by comparing the development of allergic inflammation in wildlings alongside conventional SPF mice, it is possible to test the voracity of the hygiene hypothesis, under well-controlled and standardized laboratory conditions.

In this proof-of-concept study, we set out to directly probe the hygiene hypothesis by testing whether wildlings – exposed to a rich and diverse commensal and pathogenic microbial repertoire – were protected from sensitization to common allergens and the subsequent development of allergic inflammation, when compared to SPF mice deprived of the same microbial exposure.

As published by Rosshart and colleagues, wildlings are characterized by high loads and greater diversity of bacteria, archaea, viruses, fungi as well as other eukaryotic organisms such as protozoa, mites and worms at all epithelial barrier sites including the gut, the skin and the female reproductive tract (*18*). The specific hygiene standards of wildlings and the conventional SPF mice used for this study can be found in table S1. The impact of the wildling microbiome was first analyzed on T cells from unmanipulated 8 week-old mice (fig. S1-3). Notably, the thymus of wildlings was reduced in size and cellularity (fig. S1), indicating that the wildling flora drives thymic involution. In peripheral organs including spleen, mediastinal lymph node (medLN) and lung tissue, effector (CD44^+^) CD4 and CD8 T cells were present in higher frequencies and the number of Th1, Th2 and Th17 cells were augmented in all compartments (fig. S2-3). Thus, effector T cells are more prevalent in peripheral lymphoid organs and non-lymphoid tissues of wildlings compared with SPF mice.

House dust mites (HDM) are a near ubiquitous aeroallergen to which asthmatics are commonly sensitive and which can trigger asthma exacerbations (*23–25*). In HDM-allergic asthmatics, HDM induces the production of cytokines including IL-4, IL-5 and IL-13 from Th2 cells, is bound by IgE to promote mast cell degranulation and promotes inflammation and mucus deposition in the airways (*26*). To determine if wildling mice developed allergic responses to HDM, mice were sensitized (1 μg HDM, day 0) and challenged (10 μg HDM, days 7–11) with HDM and analyzed for several parameters at day 15, the peak of airway inflammation (Fig. 1A). Histological analysis and enumeration of leukocytes in bronchoalveolar lavage (BAL), lung tissue and lung-draining medLN demonstrated a robust inflammation in the airways, lung parenchyma and medLN of wildling mice (Fig. 1B, fig. S4A). Qualitative analysis of cellular infiltrates by flow cytometry distinguished significant increases in lung tissue eosinophils, T cells, B cells, neutrophils and macrophages of wildlings administered HDM (Fig. 1C-D, fig. S1C). In addition to the enhanced inflammatory state of wildling lungs, mucus-secreting goblet cells were also more prominent in HDM-administered wildlings compared to SPF mice (Fig. 1E). HDM also induced strong serum IgG1 and IgE responses to one of the dominant allergenic proteins of HDM, Der p 2 (Fig. 1F). Hence, wildlings develop lung inflammation, goblet cell metaplasia of the airways, and systemic antibody responses at levels equal to or greater than that observed in conventional SPF mice.

**Fig. 1.**
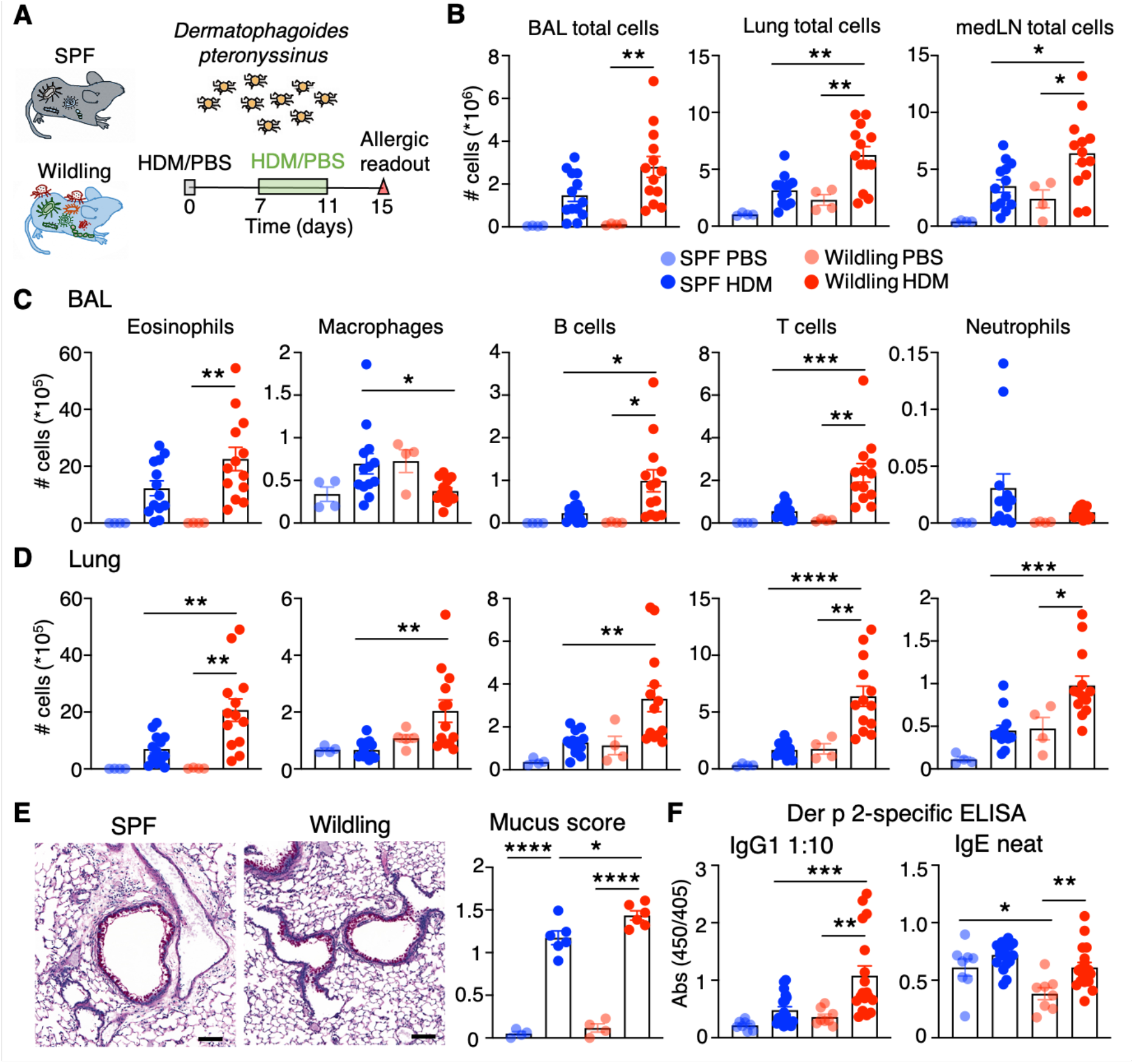
Wildling mice develop robust allergic inflammation following administration of HDM. (**A**) Regimen of HDM instillation into the airways and analysis at day 15. (**B** through **D**) Enumeration of cellular infiltrates in bronchoalveolar lavage (BAL), lung parenchyma and medLN. (**E**) Periodic Acid Schiff-Diastase (PAS-D) staining of the lungs for mucus secreting cells. Airways were scored in a blinded fashion. (**F**) Serum IgG1 and IgE specific to the Der p 2 allergen of *Dermatophagoides pteronyssinus*. One-way ANOVA and Bonferroni’s multiple comparisons test was used. In **A** through **D**, mice administered PBS n=4, mice administered HDM n=13. In **E**, PBS n=4, HDM n=6. In **F**, PBS n=8, HDM n=18-19. *P < 0.05, **P < 0.01, ***P < 0.001, ****P < 0.0001.

Heightened lung eosinophilia and mucus production indicated that Th2 cell responses may be augmented in wildling mice. T cells are activated through highly diverse T cell antigen receptors (TCRs) expressed on their cell surface, which recognize cognate peptides presented in the context of MHC molecules. Following activation by HDM-derived peptides, CD4 T cells may differentiate into Th2 cells, which express the receptor for IL-33 (comprised of subunits ST2 and IL1RAP) and secrete IL-5 and IL-13 in the lung (*27*). Th2 cell differentiation in response to HDM was measured in three ways; 1) using the non-specific mitogenic stimuli PMA and ionomycin, 2) using whole house dust mite extracts, and 3) using PE-labelled tetramers of MHC I-A^b^ loaded with Der p 1 217-227 CQIYPPNVNKI (Der p 1:I-A^b^ tetramer) (*28*). Mitogenic restimulation of leukocytes from BAL, lung tissue and medLN depicted an approximately eight-fold increase in Th2 cells producing IL-5 and/or IL-13 in wildling mouse airways, lung or medLN compared with SPF mice administered HDM (Fig. 2A-B, fig. S5A). Th2 cell numbers in wildling BAL or lung tissue far outweighed those of Th1 (IFN-*γ*^+^) or Th17 (IL-17^+^) cells, with approximately 40% of gated effector CD4 T cells in the BAL producing IL-13 (Fig. 2A-B). When leukocytes from lung or medLN were restimulated overnight with whole HDM extracts, Th2 cells producing IL-5 and IL-13 were also significantly more numerous in wildlings (Fig. 2C, fig. 5B), suggesting that HDM-specific Th2 cell responses are greater in wildling compared with SPF mice. Finally, the lung tissue of wildlings administered HDM contained similar overall numbers of Der p 1:I-A^b^ tetramer-reactive cells as SPF mice administered HDM (Fig. 2D, fig. 5C). However, Der p 1:I-A^b^ tetramer-reactive cells in wildlings were comprised of a higher frequency of Th2 cells (ST2^+^) and a reduced frequency of Treg (Foxp3^+^) compared with their SPF counterparts (Fig. 2D, fig. 5C). Of note, control wildling mice administered PBS harbored very few Der p 1:I-A^b^ tetramer-reactive cells (Fig. 2D) and did not display obvious serum antibody binding to recombinant Der p 2 (Fig. 1F, fig. S4D). This suggests that a fulminant microbial milieu does not impede, but instead amplifies the generation of *de novo* T and B cell responses to common aeroallergens, manifesting in robust allergic inflammation of the airways.

**Fig. 2.**
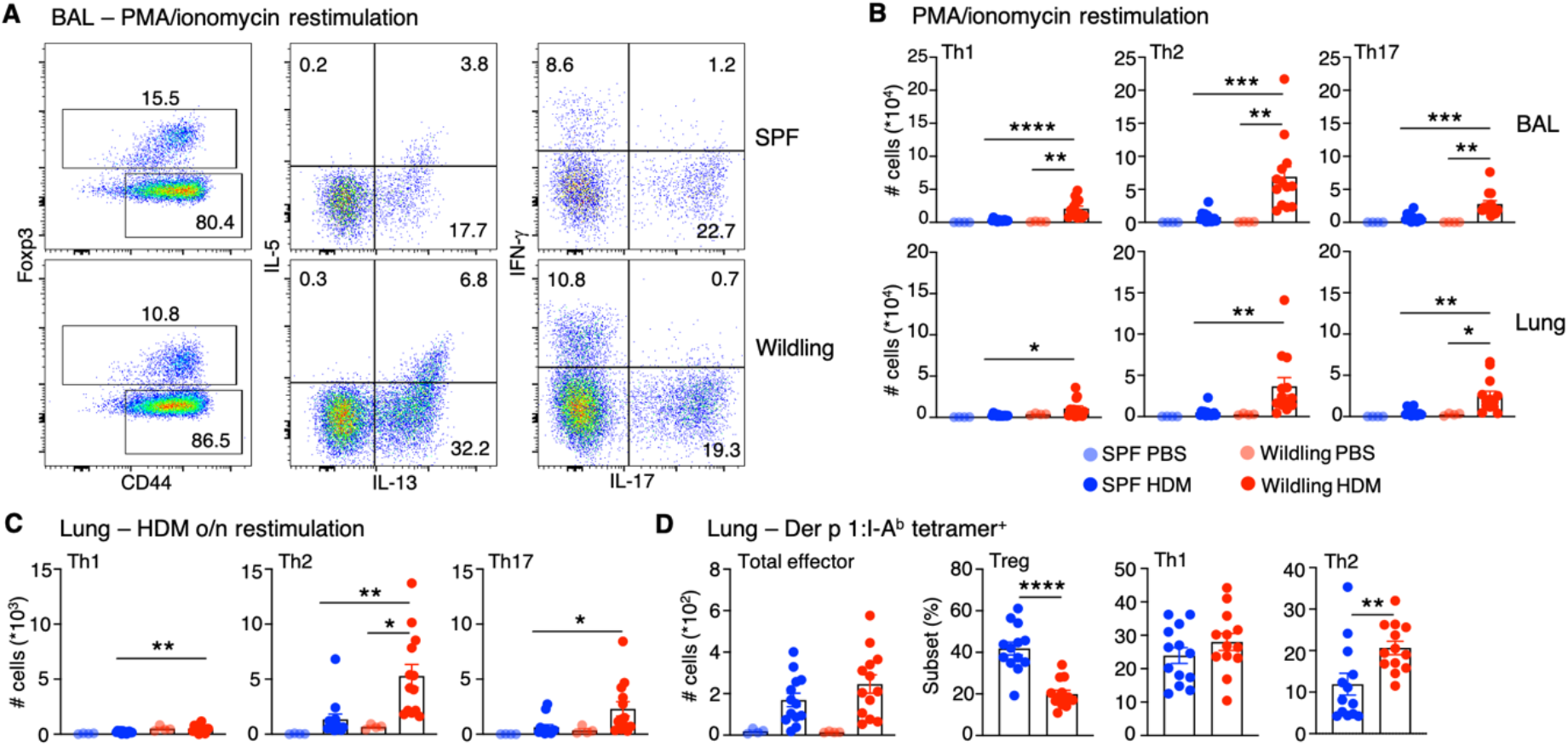
Th2 cell responses are augmented in wildlings following HDM administration. (**A**) Representative plots of CD44 versus Foxp3 (left-hand) in gated CD4 T cells, IL-13 versus IL-5 (middle) and IL-17 versus IFN-γ (right) expression in gated effector T helper cells (CD4^+^CD44^+^Foxp3^−^) following stimulation with PMA and ionomycin. (**B**) Enumeration of Th1 (IFN-γ^+^), Th2 (IL-13^+^ and/or IL-5^+^) and Th17 (IL-17^+^) T helper cells following stimulation with PMA and ionomycin. (**C**) Enumeration of Th1, Th2 and Th17 cells following overnight stimulation with HDM extracts. (**D**) Enumeration of total Der p 1:I-Ab tetramer^+^ cells and subsets therein in lung parenchyma. One-way ANOVA and Bonferroni’s multiple comparisons test was used for multiple comparisons. In **D**, Student’s t-test was used since subset frequencies were omitted from PBS mice due to low cell numbers. PBS n=4, HDM n=13. *P < 0.05, **P < 0.01, ***P < 0.001, ****P < 0.0001.

Allergic inflammation to HDM is mediated primarily by Th2 cells. Allergens with high proteolytic activity including fungal extracts are known to induce epithelial cell damage leading to the release of alarmins such as IL-33, that activate ILC2. In response to IL-33 and other innate cytokines, ILC2 expand, produce high quantities of IL-5 and IL-13 and induce allergic inflammation (*29*). The lung and bone marrow of unmanipulated wildling mice had significantly fewer ILC2 compared with SPF mice (Fig. 3A-B), suggesting that responses to alarmins may be reduced. In order to analyze the responses of SPF and wildling mice to innate stimuli, we intranasally administered mice recombinant (r) IL-33 (Fig. 3C, fig. S6) or extracts of the fungus *Alternaria alternata* (*AA*) (fig. S7) for three consecutive days. SPF and wildling mice experienced a comparably robust infiltration of eosinophils into the lung and airways at days 2 and 4 after initial airway challenge (Fig. 3C, fig. S7A). This coincided with an expansion in the lung ILC2 compartment in both SPF and wildling mice, although ILC2 numbers remained lower in wildlings throughout the time course (Fig. 3D, fig. S7).

**Fig. 3.**
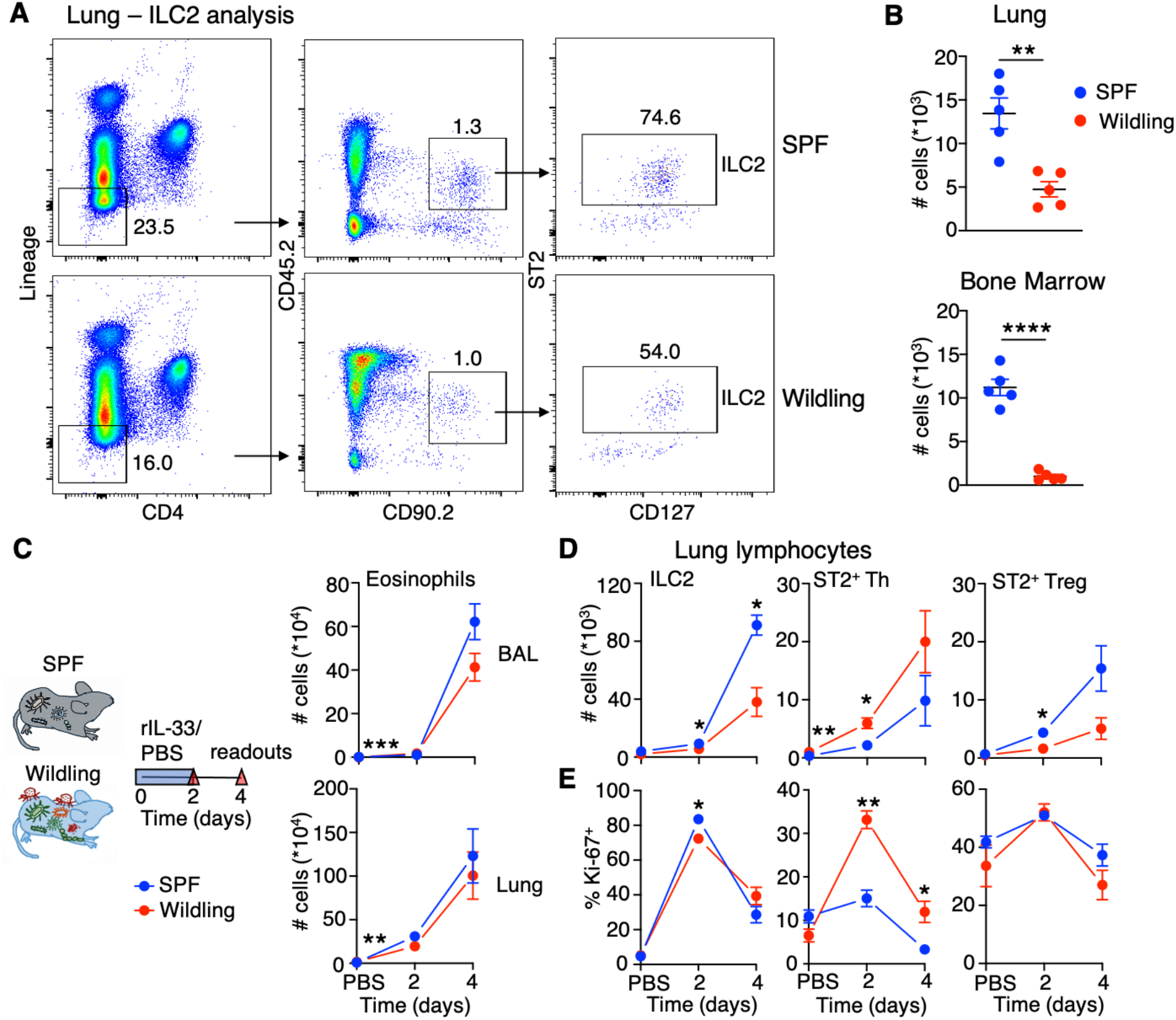
Reduced ILC2 in wildlings but robust allergic responses to rIL-33 administration. (**A**) Gating strategy for identification of ILC2. Representative plots of CD4 versus lineage (CD3, CD19, CD11c, NK1.1), CD90.2 versus CD45.2 and CD127 versus ST2. (**B**) Enumeration of lung and bone marrow ILC2. (**C**) Regimen of rIL-33 (200 ng daily for 3 consecutive days) or PBS instillation into mouse airways and subsequent analysis. Graphs of eosinophil infiltration into the airways and lung parenchyma. (**D**) Graphs of ILC2, ST2^+^ T helper and ST2^+^ Treg cell numbers in the lung following administration of rIL-33/PBS. (**E**) Graphs of the frequency of gated ILC2, ST2^+^ T helper or ST2^+^ Treg cells that were Ki-67^+^. In **A** and **B**, n=5 mice per group. In **C** through **E**, n=3 mice per group per time-point. Student’s t-test was used for all comparisons. *P < 0.05, **P < 0.01, ***P < 0.001.

The primary mode of T helper cell activation is through TCR-mediated recognition of cognate peptides presented in the context of MHC-II. However, a recent study showed that effector Th2 cells responded to innate cytokines independently of signals through the TCR (*30*). While suppressive ST2^+^ Treg outnumbered ST2^+^ Th cells in SPF mice, the reverse was true in wildings (fig. S6B). Over the short course of rIL-33 or *AA* administration in wildling mice, lung ST2^+^ Th cells expanded and maintained their majority over ST2^+^ Treg cells (Fig. 3D, fig. 6B, fig. S7A-B). ST2^+^ Th cells from wildlings rapidly entered division in response to rIL-33 (Fig. 3E) or *AA* (fig. S8), which was not observed in SPF mice. This indicates that the rich wildling microbiome primes endogenous Th2 cells to respond to alarmins triggered by incoming allergens.

Our study highlights that despite their increased microbial diversity and colonization with microorganisms previously proposed to dampen allergic sensitization such as fungi and species of *helicobacter, lactobacilli, bifidobacteria* and *bacteroides* (*10, 31–34*), wildlings generated robust allergic immune responses. If anything, the microbial exposure promoted cognate antigen-specific Th2 cell responses to HDM and non-cognate Th2 cell responses to alarmins.

Apart from microbial exposures, many lifestyle factors including sedentary and indoor living, decreased physical activity as well as unhealthy diets have been proposed to play a role in the rapid rise in allergy incidence (*35*). Moreover, amicrobial environmental factors such as the increase in air pollution and the steadily growing environmental exposure to new chemical compounds – present in nearly all common consumer products – likely contributes to the surge of allergic diseases (*36*).

This proof-of-concept study utilizes a well-controlled experimental approach and directly challenges the notion that a lack of exposure to diverse microbes is the primary factor for the rise in allergy incidence. Consequently, it emphasizes the rather multifactorial nature of allergic diseases and asks us to recalibrate our view on allergic diseases by more thoroughly considering the roles that other factors such as lifestyle, environmental pollution and new chemical compounds might play.

## Acknowledgements

The authors would like to thank the staff at KM-F animal facility for facilitating the transport and animal husbandry of wildling mice. Pathological analysis of lung tissue was performed at the Morphological Phenotype Analysis facility at Karolinska Institutet.

## Author contributions

Conceptualization: JMC.

Methodology: JM, SPR, SN, SV, HJH.

Investigation: JM, CC, JMS, ML, JMC, SN.

Data curation and visualization: JM, JMS, JMC, SPR.

Funding acquisition: JMC, SN, SPR.

Supervision: JMC, SN.

Writing — original draft: JM, JMC.

Writing — review and editing: JMC, SPR, SN.

## Supplementary Materials

### Materials and Methods

#### Mice

C57BL/6NTac murine pathogen free (MPF) mice were used in all experiments and adhered to our characterization of SPF. C57BL/6NTac wildling mice were created through inverse germ-free rederivation as described by Rosshart and colleagues (*18*). The wildling mouse colony was housed and animals were bred at the Medical Center – University of Freiburg, Germany. Wildling and C57BL/6NTac MPF mice were around 8-weeks-old at the start of experiments. Wildling mice and C57BL/6NTac MPF mice were transported at the same time to the Comparative Medicine animal facilities at Karolinska Institutet and allowed to rest for at least one week prior to the commencement of an experiment. Female mice were used in all experiments described herein. Experiments were approved by the Stockholm Regional Ethics Committee (8971/2017+B6905/2020).

#### Assessment of the hygiene standards

C57BL/6NTac wildling mice: Blood drops dried on Opti-Spot Cards (IDEXX BioAnalytics), dry fur swabs (DRYSWAB™ FLOCK from Puritan), dry oral swabs (FLOQSwabs^®^ from COPAN) and fecal pellets were collected according to the manufacturer’s sampling guidelines and screened for microorganisms with two independent methods (PCR and serology) utilizing the “Mouse 3R Comprehensive Serology Panel” as well as the “Mouse 3R Quarantine Annual SOPF PCR Panel” from IDEXX BioAnalytics. The assays were performed on pooled samples, a microorganism was considered present, if it was identified through at least one of the two independent methodologies. Conventional specific-pathogen-free C57BL/6NTac mice: The hygiene standard was assessed and reported by the mouse vendor (Taconic Biosciences).

#### Models

##### House dust mite (*Dermatophagoides pteronyssinus*)

Eight-week-old mice were anesthetized with isoflurane and sensitized with 1 μg of HDM (Stallergenes Greer) in 40 μl of PBS intranasally. Seven days after sensitization, mice were challenged with 10 μg of HDM for 5 consecutive days. Mice were sacrificed and organs harvested 4 days after the last challenge. BAL was performed by three consecutive flushes of the airway with 1 ml of PBS.

##### Recombinant IL-33 (rIL-33) or *Alternaria alternata* (*AA*) extract

rIL-33 (200 ng) or *AA* extracts (20 μg) were administered intranasally for three consecutive days. Mouse BAL, lung tissue and medLN was harvested on day 2 (one day after the second dose of rIL-33 or *AA*) and on day 4 (2 days after the last dose of rIL-33 or *AA*).

#### Organ processing

The lungs, medLN, mesLN, thymus and spleen were pushed through 100 μM sieves in around 10 ml volume of 2% heat-inactivated FCS (Sigma) in PBS (FACS buffer) to achieve single cell suspensions. Femurs were flushed with cold FACS buffer using 23-25G needles. Total lung, spleen and bone marrow single cells were lysed of red blood cells using a hypotonic buffer. Cells were resuspended in either FACS buffer or tissue culture medium, depending on the requirement for restimulation.

#### Restimulation of T cells

To detect cytokine production, cells were cultured in IMDM supplemented with penicillin/streptomycin, glutamine, 2-mercaptoethanol (all from Invitrogen), and 10% heat-inactivated FCS (Sigma). For restimulation with PMA (50 ng/ml) and ionomycin (5 μM), Brefeldin A (Sigma) was added from the beginning of culture and cells were harvested 3 hours later. For HDM-specific restimulation, cell suspensions were plated in 48-well plate with 20 μg/ml HDM extracts. After 6 hours, Brefeldin A was added to culture and after an additional 9-hour culture, cells were collected and stained with antibodies for FACS analysis. The eBioscience Fixation Kit was used when intranuclear staining of Foxp3 was performed.

#### Derp1 217-227 CQIYPPNVNKI I-A(b) Tetramer staining

I-A(b) house dust mite Der p 1 217-227 CQIYPPNVNKI was ordered from NIH Tetramer Core Facility. Half of the lung cells were resuspended in 250 μl FACS buffer and Fc block, rat and mouse serum (all in 1/100 dilution) were added, mixed well and incubated for 10 mins at room temperature (RT). Then, the total volume was topped to 500 μl with FACS buffer and 5 μl PE-labeled tetramer was added for 1h at RT. After incubation, EasySep™ Mouse PE Positive Selection Kit II (STEMCELL, Catalog #17666) was used to select PE-labeled tetramer cells. Briefly, cells were washed once with 10ml cold FACS buffer, and resuspended in 250 μl MACS buffer (0.5% FBS, 2 mM EDTA in PBS) with 25 μl PE Selection Cocktail and incubated for 15 minutes at RT. Thereafter, 15 μl Dextran RapidSpheres™ was added and incubated for additional 10 mins at RT. Cells were then washed and selected with magnets according to manufacturer’s protocol.

#### Flow cytometry

Flow cytometry was performed on a BD LSRII with combinations of the following antibodies: BD: CD19 (1D3), CD11c (HL3), B220 (RA3-6B2), CD3 (145-2C11), CD4 (RM4-5 and GK1.5), CD8 (53-6.7), CD44 169(IM7), GR-1 (RB6-8C5), IFN-γ (XMG1.2), IL-4 (11B11), IL-17 (TC11-18H10), Ki-67 (B56), CXCR3 (CXCR3-173), CD90.2 (53-2.1), CD45.2 (104), CD25 (PC 61), CD69 (H1.2F3), CD49d (R1-2), Siglec-F (E50-1702440) and Purified Rat Anti-Mouse CD16/CD32 (2.4G2); from Invitrogen: CD127 (A7R34), CD11c (N418), FOXP3 (FJK-16s), IL-13 (ebio13A); from Biolegend: IL-5 (TRFK5), NK1.1 (PK136), ST2 (DIH9). Fixable viability dye eFluor 780 from Invitrogen. All samples were fixed before being run on the flow cytometer.

#### Histopathological analysis

Lungs were inflated with and fixed in 10% Formalin for a minimum of 24 hours before being embedded in paraffin. 4 μM sections were cut on a rotary microtome (Mikrom HM355s) and periodic acid-Schiff-diastase (PAS-D) and Hematoxylin & Eosin (H&E) stains were performed. For analysis of mucus score, PAS-D-stained sections were scored on a 0-4 point scale with points assigned based on the percentage of the airway covered by positively stained cells; 0 points for 0% of the airway affected, 1 point for 1-25%, 2 points for 26-50%, 3 points for 51-75% and 4 points for more than 75% PAS positive. A minimum of 38 full airways were counted per mouse lung. For analysis of epithelial thickness; 1 point was assigned for airways where the epithelium was not a monolayer and had >1 cell. 1 point for the presence of clusters of airway epithelial cells. For assessment of inflammation around the airways; 0 points for no inflammation, 1 point for some inflammatory cells around airway, 2 points for a ring of inflammatory cells around the airway, 3 points for a ring 2-4 cells deep and 4 points for a ring more than 4 cells deep.

#### ELISA

For HDM experiments, allergen extracts (5 μg/ml) or recombinant Der p 2 (3 μg/ml) was diluted in PBS and used to coat ELISA plates (nunc). Plates were incubated at 4°C overnight and blocked with 2% milk in PBS the following morning. Serum was diluted 1:10 for detection of IgG1. For detection of allergen-specific IgE, serum was depleted of IgG first using Protein G HP SpinTrap™ columns according to the manufacturer’s protocol. Plates were washed in PBS/Tween. Serum was added to the wells and incubated for 2 h at room temperature. Plates were washed again and incubated for one hour with secondary antibody, either HRP coupled anti-IgG1 (Southern Biotech, 1070-05) or biotin coupled anti-IgE (BD, R35-72) followed by streptavidin – HRP (Mabtech). TMB substrate (KPL) followed by H_2_SO4 were used to develop and stop the reaction. The Asys Expert 96 ELISA reader (Biochrom) was used to read OD at 450 nm/405nm.

#### Statistical analysis

The statistical analysis used is stipulated in each figure. Student’s t test was used to compare two groups. One-way ANOVA and Bonferroni’s multiple comparisons test was used when more than two comparisons were made. Mean and SEM are shown in all graphs. *P < 0.05, **P < 0.01, ***P < 0.001, ****P < 0.0001.

### Supplementary text

#### Wildling mice are colonized by a diverse range of commensals and pathogens

The hygiene standards of wildling mice were determined using two methods, the Mouse 3R Comprehensive serology panel and the mouse 3R Quarantine Annual SOPF PCR panel. These are reported in table S1.

Several studies have shown an inverse correlation in microbial diversity and the risk of developing allergy (*34, 37, 38*). Wildling mice have considerably greater diversity in the repertoire of microbes in their housing, and in the microbiome of the gut, skin and vagina (reported in (*18*)). Colonization with families of gram-negative bacteria including *Campylobacteraceae, Neisseriaceae, Helicobacteraceae* and *Pasteurellaceae* is a feature of wildling skin. As regards specific commensals proposed to regulate the development of allergy including *lactobacilli* and *bacteroides*, wildlings harbor increased levels of these genera and increased levels of the phylum *Actinobacteria*, which includes *bifidobacteria spp* (*33, 37, 39-45*). Fungi (in particular of *Ascomycota*) and pinworms are also prominent in wildlings, which may play a role in allergic sensitization (*31, 34, 46, 47*). In all, wildlings are marked by considerably greater microbial diversity than SPF mice, including with several microbes proposed to regulate allergic sensitization and disease.

#### Wildlings are characterized by reduced thymus cellularity

We characterized the immune cell composition of organs including the thymus, spleen, lung and mediastinal lymph nodes. The thymus of wildling mice was found to contain fewer total lymphocytes compared to SPF mice. CD4^+^CD8^+^ (DP) thymocytes, mature CD4 T cells (CD4^+^CD8^−^) and mature CD8 T cells (CD8^+^CD4^−^) were all reduced in quantity (fig. S1A-B). CD4^−^CD8^−^ (DN) thymocytes were inflated in proportion but appeared to be comprised of standard frequencies of DN1-DN4 subsets (fig. S1C-D).

#### Antigen-experienced CD4 and CD8 T cells are present in peripheral lymphoid and non-lymphoid organs of wildling mice

SPF and wildling mice had comparable numbers of total CD4 and CD8 T cells in the spleen and lung. However, CD44^+^ effector CD4 and CD8 T cells were highly enriched in the spleen and lungs of wildling mice (fig. S2). Foxp3^+^ suppressive Treg cells appeared reduced in frequency, as previously published (*18*), but were numerically similar between SPF and wildling spleen (fig. S2). The effector CD8 T cell population of wildling mouse spleen and lung was enriched for ‘true memory’ CD8 T cells (CD8T™) while the numbers of ‘virtual memory’ CD8 T cells (CD8T_VM_) (*48*) were comparable in spleen, and increased in the lung (fig. S2). The mediastinal lymph node (medLN) was larger in wildlings compared to SPF mice and all cell populations characterized were increased, including naïve cell populations (fig. S2). To characterize cytokine production by T cells, leukocyte preparations from SPF and wildling mice were stimulated with phorbol-12-myristate-13-acetate (PMA) and ionomycin for 3 hours. Effector T helper cells (CD3^+^CD4^+^CD44^+^Foxp3^−^) expressing the cytokines IFN-γ (Th1), IL-5 and/or IL-13 (Th2), or IL-17 (Th17) were observed at comparable frequencies but present in significantly higher numbers in wildling spleen, lung and mediastinal lymph node compared to SPF mice (fig. S3). In the CD8 T cell compartment, wildling mice had a higher frequency and overall number of IFN-γ-producing cells (fig. S3). Thus, wildling mice are marked by diminished thymus cellularity and a greater abundance of effector CD4 and CD8 T cells in peripheral lymphoid and non-lymphoid organs.

#### Inflammatory features of mice administered HDM extracts

Administration of HDM into wildling mice induced robust signs of inflammation in the lung parenchyma, airways and circulation (Fig. 1). To decipher the quality of cellular inflammation in mice administered HDM (fig. S4A), lungs were inflated with formalin and after fixation, were sectioned for hematoxylin and eosin staining and Periodic acid-Schiff diastase staining. HDM administration into SPF or wildlings induced a marked inflammation surrounding the airways, with signs of epithelial thickening in mice administered HDM (fig. S4B). Overall, lung sections from wildling and SPF mice showed a similar level of inflammation. Airway and lung parenchyma cells were stained with markers allowing for discrimination of eosinophils, neutrophils, alveolar macrophages, T cells and B cells (reported in Fig. 1, gating strategy shown in fig. S4C), which demonstrated a robust inflammation in wildling lungs that had been challenged with HDM.

#### Th2 cell responses to HDM are heightened in wildlings compared with SPF mice

To complement the analysis of T helper cell cytokine production in lung and airways presented in Fig. 2 (main text), lymphocytes from medLN were also restimulated with PMA/ionomycin briefly or overnight with HDM. Following PMA/ionomycin stimulation, wildling mice administered HDM contained significantly higher numbers of Th1, Th2 and Th17 cells compared with SPF controls (fig. S5A). After overnight culture in the presence of HDM, only Th2 cells were present in greater quantity in wildlings, suggesting that HDM induced a more potent Th2 cell response in wildlings than in SPF mice (fig. S5A). Some IL-4 production was also detected in IL-13^+^ effector Th cells restimulated in the presence of HDM, although the magnitude of response did not differ significantly between SPF and wildling mice (fig. S5B).

To analyze the antigen-specific response to HDM in a more sensitive manner, lung cells were incubated with PE-labelled tetramers of I-A^b^ loaded with Der p 1 217-227 CQIYPPNVNKI (Der p 1:I-A^b^ tetramer). This was previously identified as a dominant response to HDM in B6 mice (*28, 49*). The results reported in Fig. 2D of the main text are based on staining with Foxp3, CXCR3 and ST2 within Der p 1:I-A^b^ tetramer-reactive cells (fig. S5C). Wildling mice administered PBS had very few effector CD4 T cells that bound the tetramer, suggesting no prior exposure to HDM (*D. pteronyssinus*) (Fig. 2D). In mice exposed to HDM, Treg, Th1 or Th2 cells could be clearly demarcated within Der p 1:I-A^b^ tetramer-reactive cells (fig. S5C). The frequency of Treg cells was reduced while the frequency of Th2 cells was increased in wildling compared with SPF Der p 1:I-A^b^ tetramer-reactive cells.

#### ST2^+^ T helper cells are significantly more numerous in wildling mice and rapidly respond to inflammatory danger signals

In SPF mice, approximately 10% of suppressive Treg (Foxp3^+^) cells express the IL-33R subunit, ST2. Comparatively fewer conventional T helper (Foxp3^−^) cells express ST2, in line with the relative paucity of Th2 cells in clean SPF mice. We noted that wildling lungs contained considerably more ST2^+^ T helper cells (fig. S6A), while the proportion of ST2^+^ Treg cells was not significantly different. Several recent reports have ascertained that ST2^+^ Treg in visceral adipose tissue, muscle and lungs are highly responsive to rIL-33 administration and may help to constrain inflammation in these tissues (*50–53*). We noted that the ratio of ST2^+^ T helper cells to ST2^+^ Treg in wildlings was greater than 2:1, while in SPF mice, ST2^+^ Treg far outnumbered ST2^+^ T helper cells (fig. S6B). Over the course of rIL-33 administration in wildlings, the ratio of ST2^+^ T helper cells to ST2^+^ Treg was maintained and even slightly increased, suggesting that ST2^+^ T helper cells responded to rIL-33 without requiring exogenous instillation of peptide antigens to stimulate the TCR.

Instillation of wildling and SPF mice with *AA* extracts also led to a rapid inflammatory response in the lung and airways. Although generally less potent than rIL-33, *AA* induced airway and lung eosinophilia, and neutrophilia in wildling mice, to a similar degree as in SPF mice (fig. S7A). ILC2 were reduced in wildlings administered *AA* throughout the time-course while ST2^+^ T helper cells were higher at day 2 in wildling administered *AA*, but not at day 4 (fig. S7B). The ratio of ST2^+^ T helper cells to ST2^+^ Treg was maintained at 2:1 in wildlings and increased in SPF mice over 4 days, suggesting that *de novo* T helper cell responses were generated in SPF mice (fig. S7C).

The expansion of ST2^+^ T helper cells and ST2^+^ Treg cells following rIL-33 and *AA* administration in these short-term instillation models prompted us to analyze Ki-67 expression, which marks cells in division. In both models, ILC2 rapidly entered division in both wildling and SPF mice (Fig. 3E, fig. S8B). A significant rise in the proportion of Ki-67^+^ cells was also observed specifically in wildling ST2^+^ T helper, indicating that endogenous Th2 cells in wildlings are capable of dividing and expanding in response to innate cytokines in the absence of the instillation of exogenous cognate antigens (Fig. 3E, fig. S8A shows the gating strategy).

No such spike in division was observed in ST2^+^ T helper cells from SPF mice. A similar response was observed following AA challenge, where a significant increase in Ki-67^+^ ST2^+^ T helper cells was observed 2 days after the first allergen administration (fig. S8B). Thus, ST2^+^ T helper cells appear to respond rapidly to innate inflammatory signals in wildlings, but not in SPF mice.

---

**Fig. S1.**
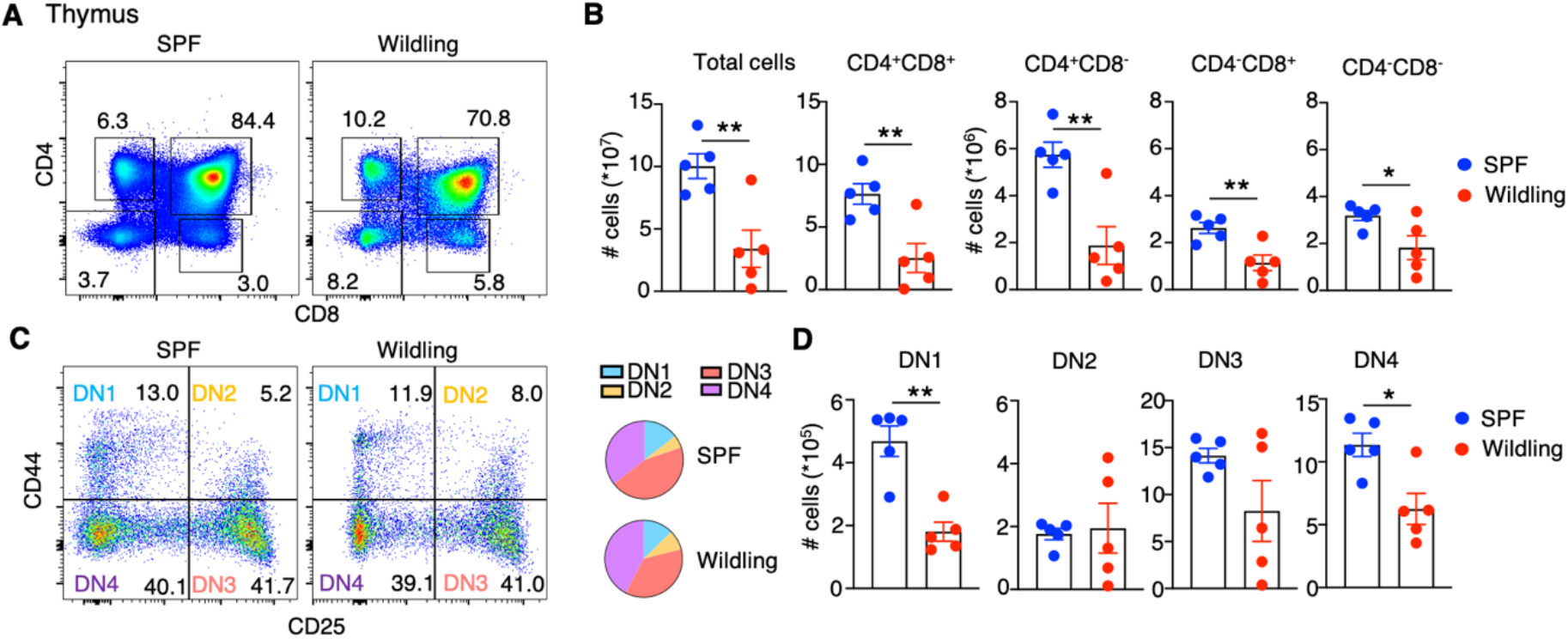
T cells of an activated phenotype are present in high numbers in wildling mice. (**A**) Representative plots of CD8 versus CD4 expression in thymus of SPF and wildling mice. Numbers indicate frequency of cells in the indicated gate. (**B**) Bar graphs depict the total cells and number of CD4^+^CD8^+^, CD4^+^CD8^−^, CD4^−^CD8^+^, CD4^−^CD8^−^ cells in the thymus of SPF (blue) and wildling (red) mice. (**C**) Representative plots of CD25 versus CD44 expression on double negative (CD4^−^CD8^−^) cells in the thymus of SPF and wildling mice. Pie charts summarizing the distribution of different developmental stages. (**D**) Bar graphs depict the total number of DN1-DN4 cells in SPF (blue) and wildling (red) mice. Student’s t-test was used for all comparisons. Each dot represents one mouse, n=5 mice per group, means and SEM are depicted, *P < 0.05, **P < 0.01.

**Fig. S2.**
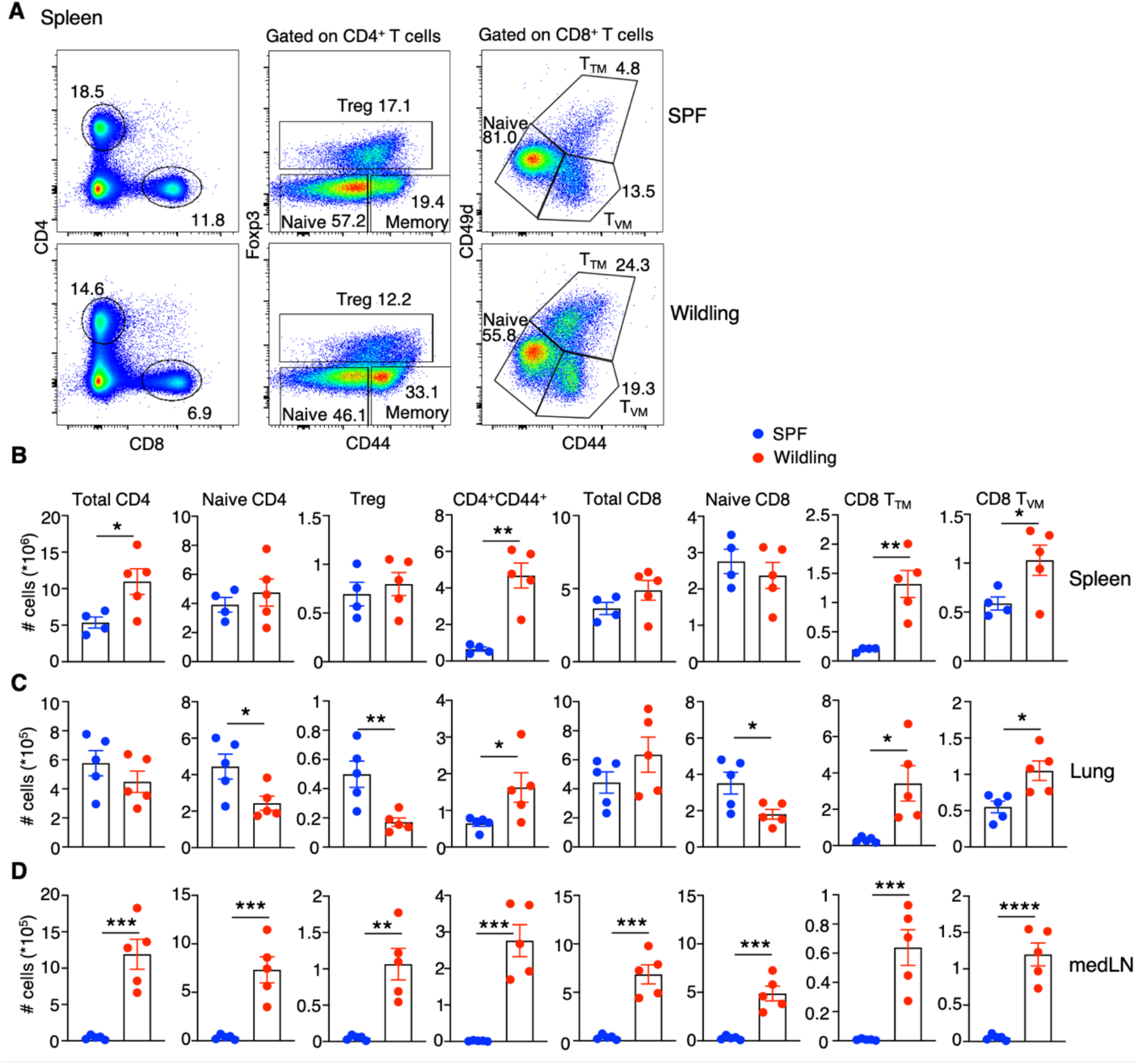
An inflated pool of antigen-experienced T cells resides in peripheral organs of wildling mice. Spleen, lung and medLN was harvested from SPF and wildling mice. (**A**) Left: representative plots of CD8 versus CD4 expression in the spleen of SPF and wildling mice, numbers indicate percentages in the drawn gates. Middle: representative plots showing the expression of CD44 versus Foxp3 within gated CD3^+^CD4^+^ T cell populations. Right: the expression of CD44 and CD49d within gated CD3^+^CD8^+^ T cell populations of SPF and wildling spleen. (**B** through **D**) Bar graph quantification of CD4, CD8 T cells in spleens (**B**), Lung (**C**), and medLN (**D**) of SPF and wildling mice, and CD4 naïve (CD3^+^CD4^+^CD44^−^), effector (CD3^+^CD4^+^CD44^+^Foxp3^−^), Treg (CD3^+^CD4^+^Foxp3^+^), CD8 naïve (CD3^+^CD8^+^CD44^−^CD49d^−^), T_VM_ (CD3^+^CD8^+^CD44^+^CD49d^−^), and T™ (CD3^+^CD8^+^CD44^+^CD49d^+^) cells. Student’s t-test was used for all comparisons. Each dot represents one mouse, n=5 mice per group, means and SEM are depicted, *P < 0.05, **P < 0.01, ***P < 0.001.

**Fig. S3.**
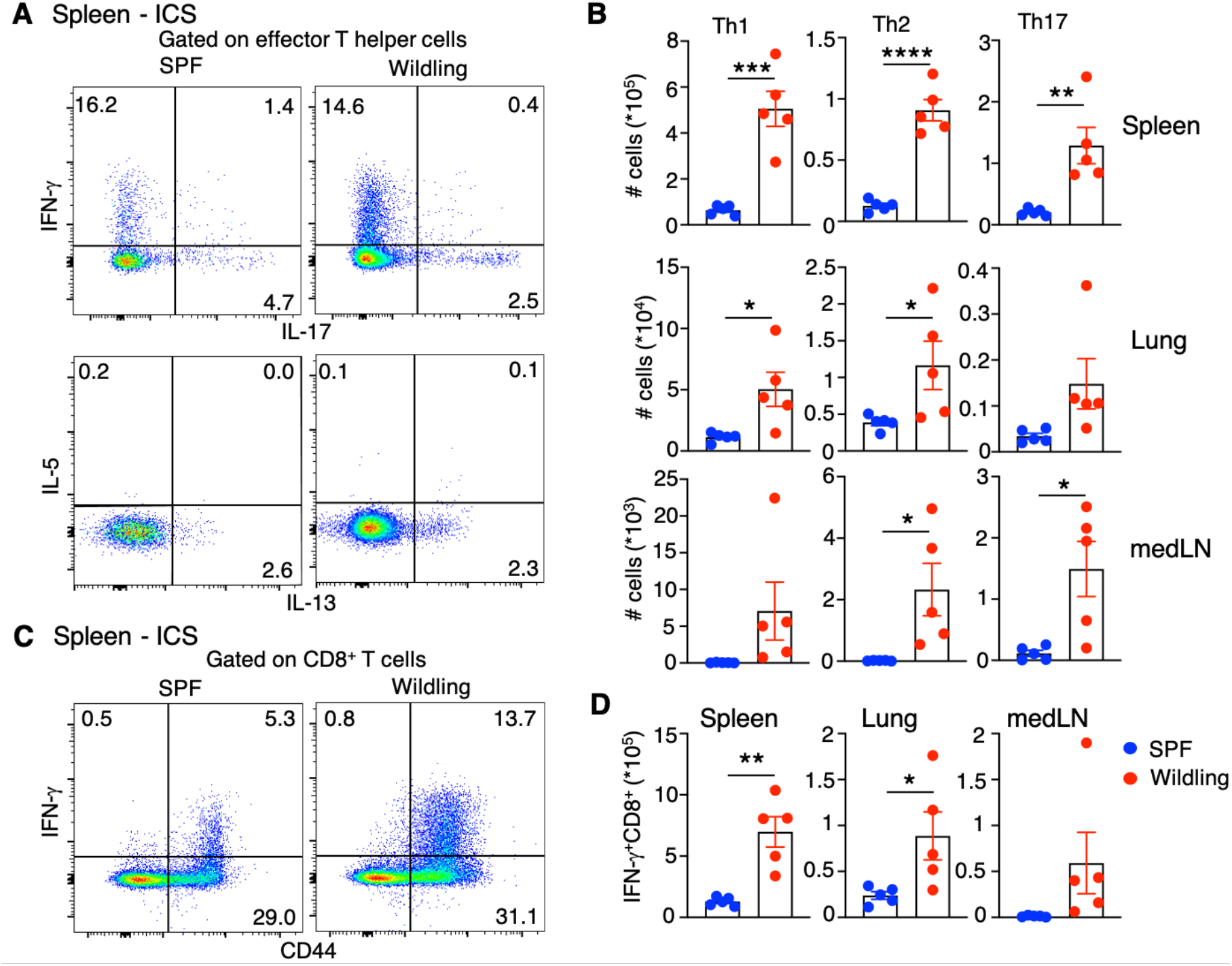
Wildlings harbour more cytokine-producing T cell subsets in peripheral organs. (**A**) Representative plots of IL-17 versus IFN-γ and IL-13 versus IL-5 expression on gated effector CD44^+^CD4^+^Foxp3^−^ T cells from the spleen. (**B**) Bar graph quantification of Th1 (IFN-γ^+^), Th2 (IL-5^+^ and/or IL-13^+^), Th17 (IL-17^+^) cell populations in spleen, lung and mediastinal lymph nodes. (**C**) Representative plots of CD44 versus IFN-γ expression on gated CD8 T cells from the spleen. (**D**) Number of IFN-γ^+^ CD8^+^ cells in spleen, lung, and medLN of SPF and wildling mice after stimulation for 3 hours in the presence of PMA and ionomycin. n=5, Student’s t-test was used for all comparisons. *P < 0.05, **P < 0.01.

**Fig. S4.**
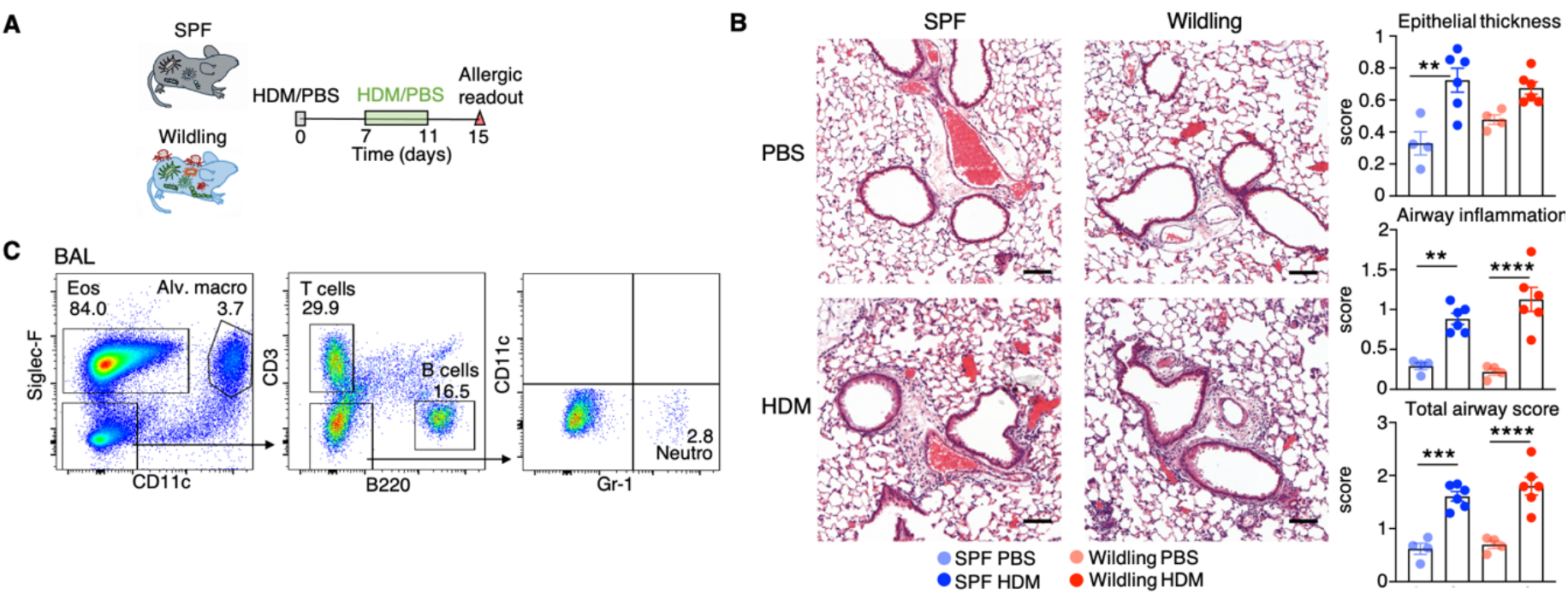
Wildling mice develop robust pathogenic responses to HDM. (**A**) Schematic of intranasal HDM instillations. Sensitization with 1 μg HDM and challenge with 10 μg of HDM. (**B**) Representative Hematoxylin & Eosin (H&E)-stained lung tissue from SPF or wildling mice treated with PBS/HDM, scale bars depict 100 μM. Epithelial thickness, airway inflammation and total (epithelium + airway inflammation) scores were calculated. Slides were scored in a blinded fashion. PBS n=4, HDM n=6. (**C**) Representative plots of BAL showing the gating strategy of eosinophils (Siglec-F^+^CD11c^−^), alveolar macrophages (Siglec-F^+^CD11c^+^), T cells (Siglec-F^−^CD11c^−^B220^−^CD3^+^), B cells (Siglec-F^−^CD11c^−^CD3^−^B220^+^), and neutrophils (Siglec-F^−^CD11c^−^B220^−^CD3^−^Gr-1^+^). One-way ANOVA and Bonferroni’s multiple comparisons test was used for all 4 comparisons. **P < 0.01, ***P < 0.001, ****P < 0.0001.

**Fig. S5.**
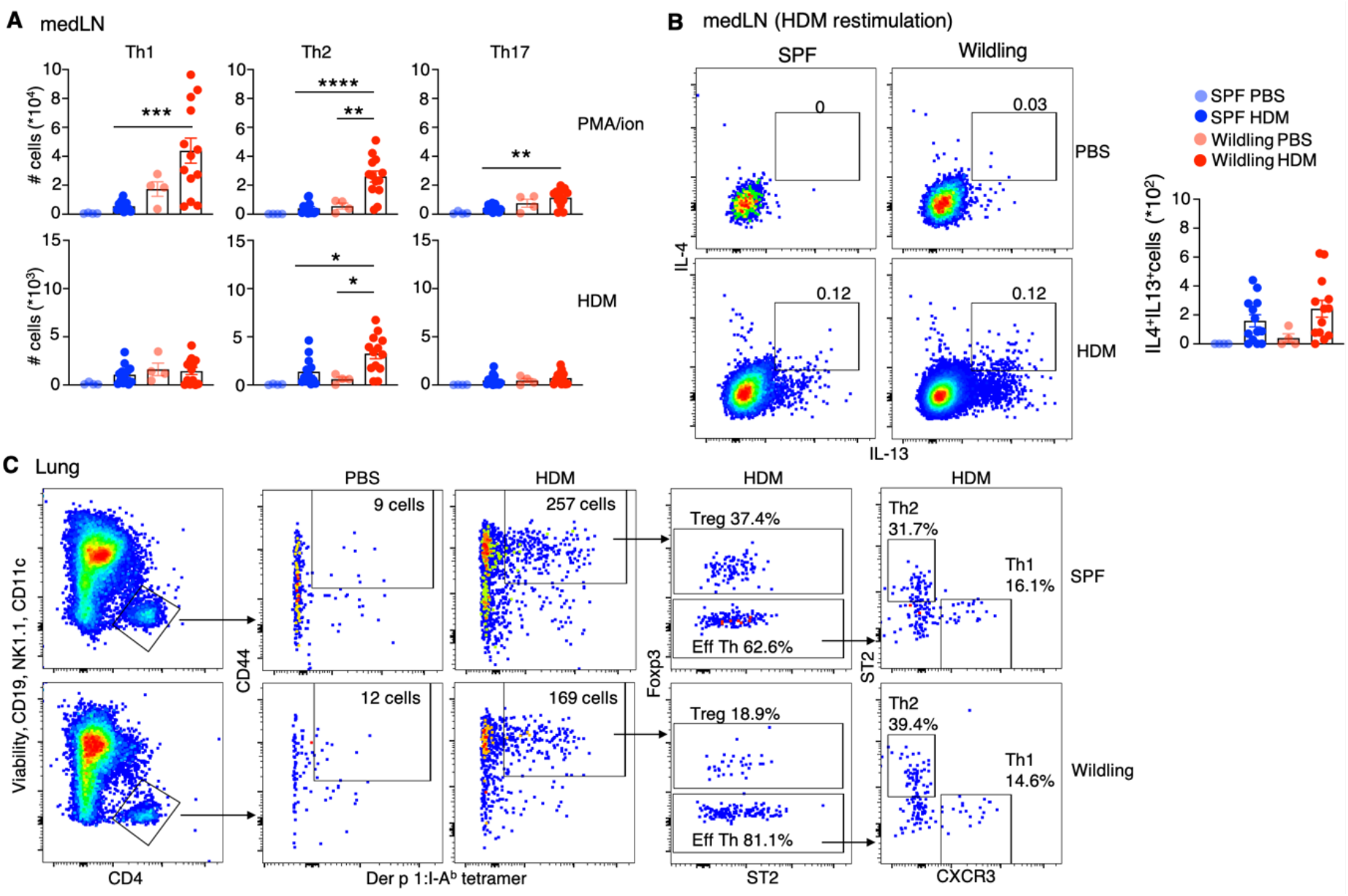
Characterising the T helper cell response to HDM. (**A**) Absolute numbers of Th1, Th2 and Th17 cells following PMA and ionomycin or overnight HDM stimulation of medLN. (**B**) Representative plots of IL-13 versus IL-4 expression on HDM-restimulated medLN and the corresponding graph of the number of IL-4^+^IL-13^+^ CD4 T cells. (**C**) Gating scheme to identify effector (CD44^+^) Der p 1:I-A^b^ tetramer-reactive in the lung of SPF and wildling mice. In main figure 2D, the frequency of Treg, Th1 and Th2 cells among Der p 1:I-A^b^ tetramer^+^ CD44^+^ CD4^+^ T cells is shown. One-way ANOVA and Bonferroni’s multiple comparisons test was used for all 4 comparisons. PBS n=4, HDM n=13. *P < 0.05, **P < 0.01, ***P < 0.001, ****P < 0.0001.

**Fig. S6.**
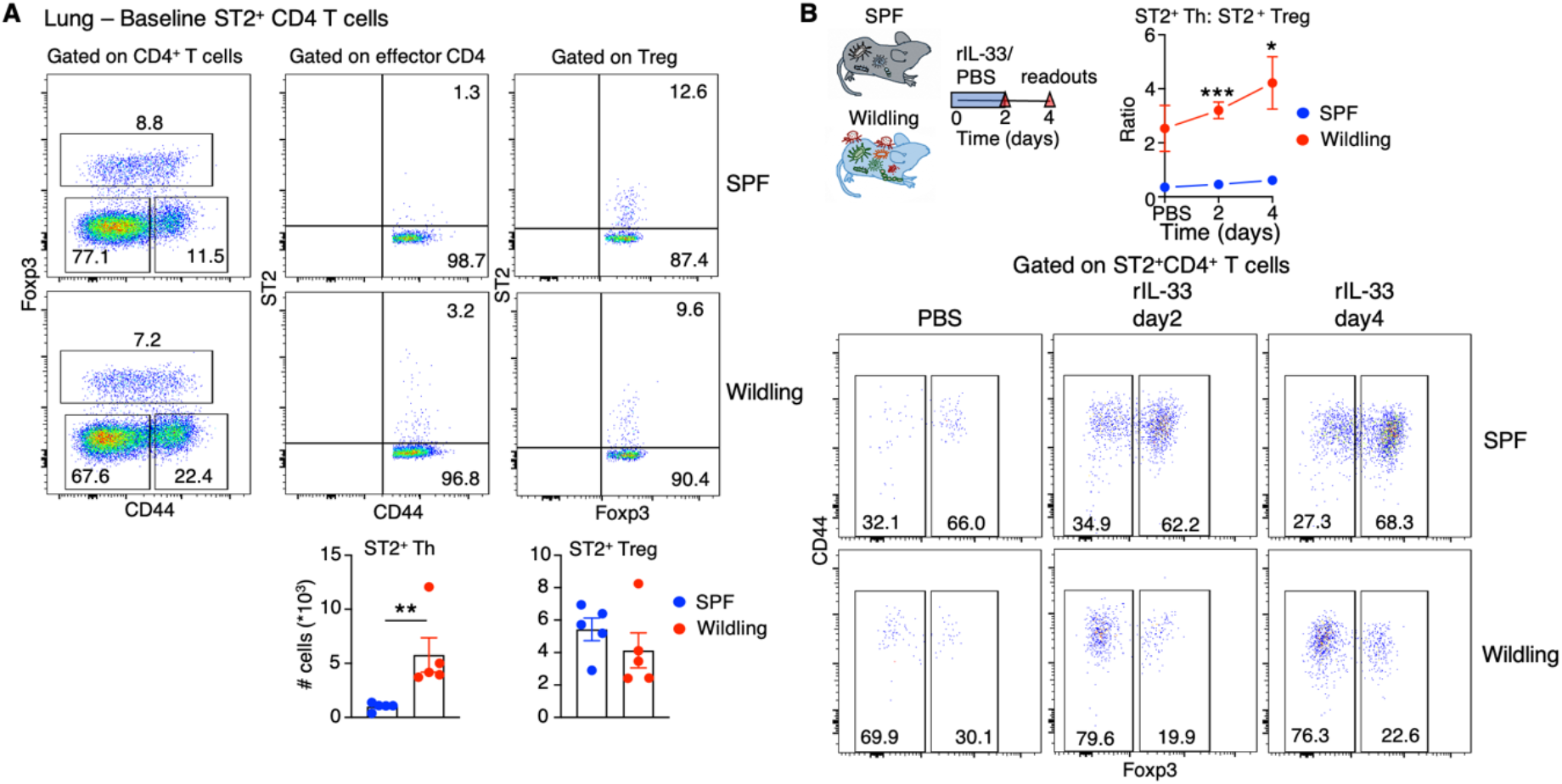
The number of ST2^+^ effector T helper cells is higher in wildling compared with SPF mice. (**A**) Representative plots of CD44 versus Foxp3 expression on gated CD4 T cells in the lung (Left-hand plot). Representative plot of CD44 versus ST2 on gated effector CD44^+^Foxp3^−^ CD4 T cells (middle plot), and Foxp3 versus ST2 on gated Treg (right-hand plot). Graphs of ST2^+^ effector CD4 T and ST2^+^ Treg are shown below, n=5 mice per group (**B**) Schematic of intranasal rIL-33 (200 ng daily for 3 consecutive days) instillations and ratio of ST2^+^ Th:ST2^+^ Treg across the time course (top). Representative plots of Foxp3 versus CD44 expression within gated ST2^+^CD4^+^ T cells, n=3 per group per time-point. Student’s t-test was used for all comparisons. *P < 0.05, ** P<0.01, ***P < 0.001.

**Fig. S7.**
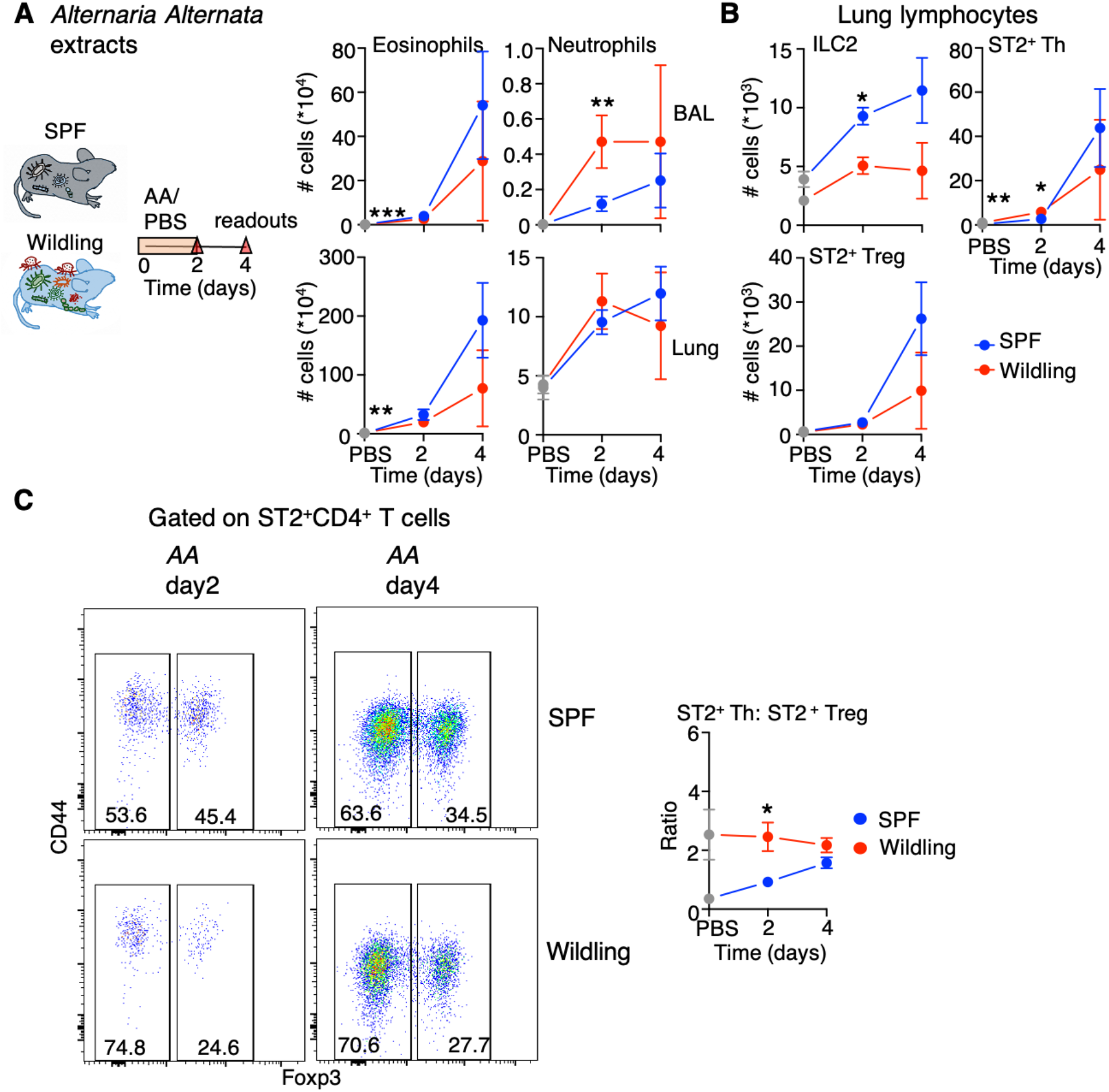
Wildling mice develop allergic responses to *Alternaria alternata* (*AA*) extracts. (**A**) Schematic of intranasal *AA* (20 μg daily for 3 consecutive days) instillations and absolute numbers of eosinophils and neutrophils in BAL and Lung. (**B**) Absolute numbers of ILC2, ST2^+^ effector (CD44^+^) CD4 T and ST2^+^ Treg in lung of SPF and wildling mice after *AA* administration. (**C**) Representative plots of Foxp3 versus CD44 expression within gated ST2^+^CD4^+^ T cells and the ratio of ST2^+^ Th:ST2^+^ Treg after *AA* administration. PBS, *AA* and rIL-33 instillations were conducted as part of the same experiment. PBS symbols are shown in gray to denote that these are the same control values shown in the rIL-33 graphs. n=3 per group per time-point. Student’s t-test was used for all comparisons. *P < 0.05, ** P < 0.01, ***P < 0.001.

**Fig. S8.**
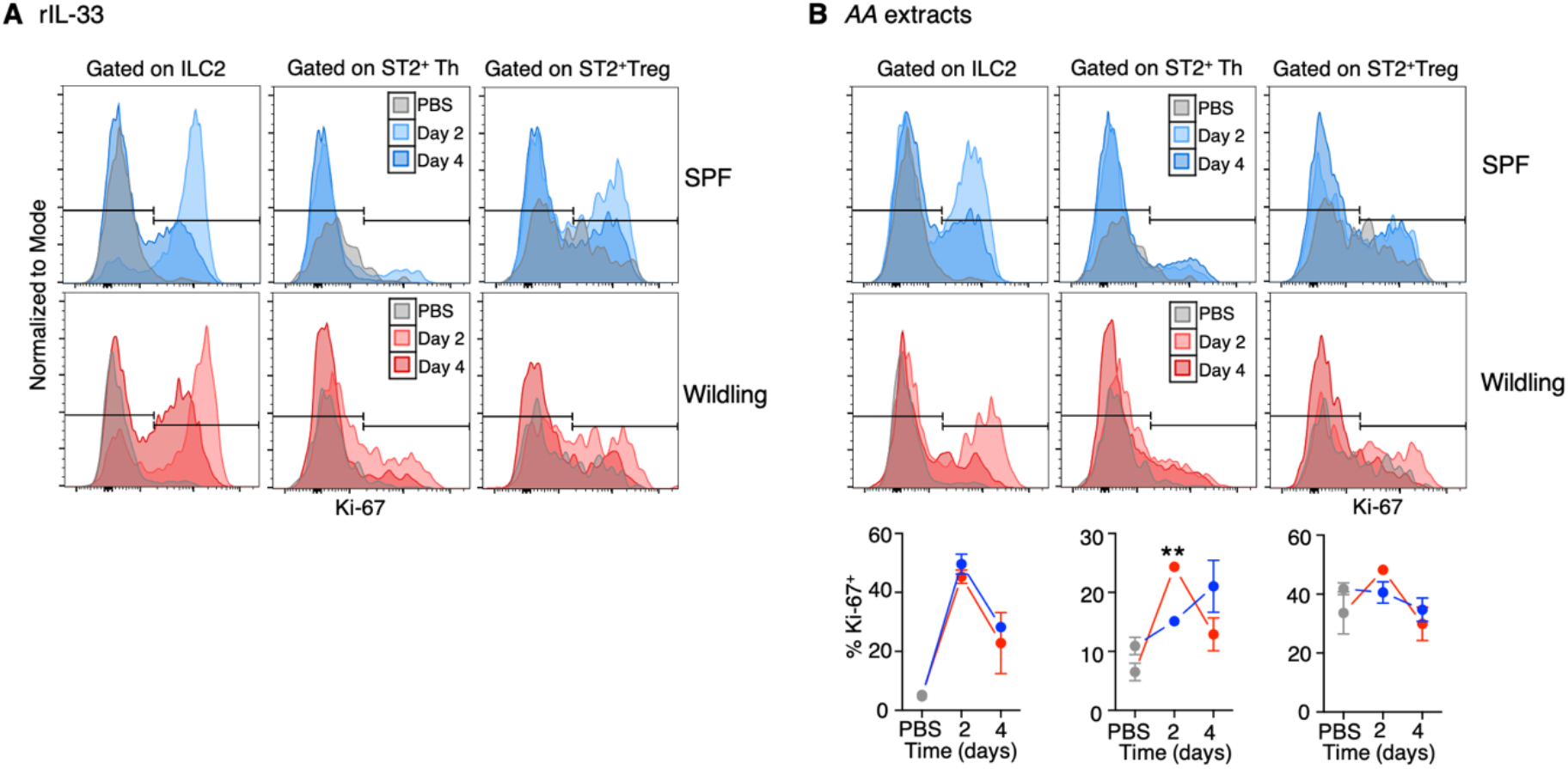
ST2^+^ Th cells in wildling mice rapidly enter division after rIL-33 or *AA* instillations. (**A** and **B)** Histograms showing the expression of Ki-67 within gated ILC2, ST2^+^ effector Th and ST2^+^ Treg after rIL-33 (**A**) or *AA* extract instillations (**B**). Ki-67 quantification for rIL-33 is shown in Fig. 3E of the main article. Quantification of Ki-67 in the indicated subset is shown for *AA* instillation (**B**), n=3 per group per time-point. Student’s t-test was used for all comparisons. **P < 0.01.

**Table S1.**
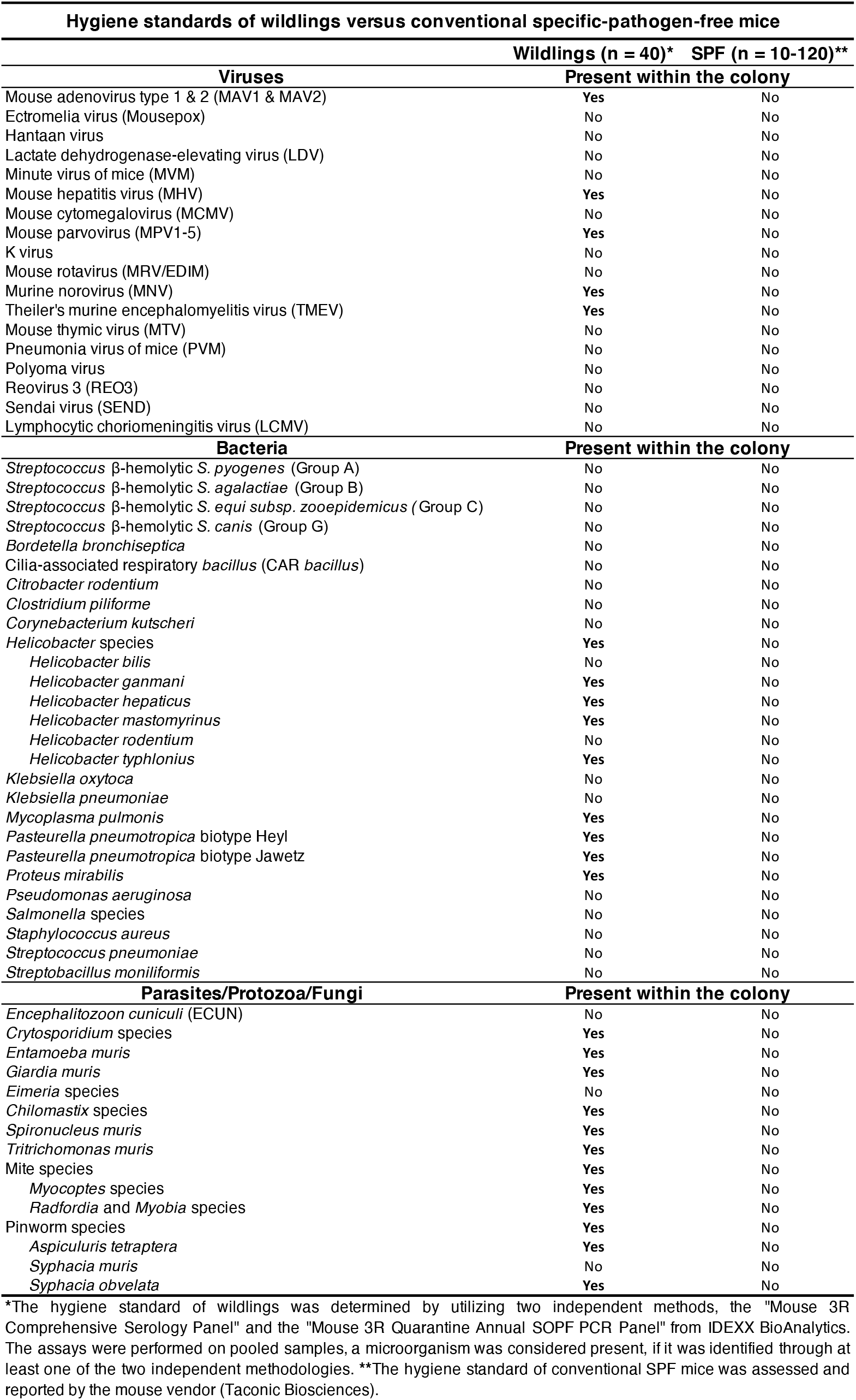
Hygiene standards of wildlings versus conventional specific-pathogen-free mice. *The hygiene standard of wildlings was determined by utilizing two independent methods, the “Mouse 3R Comprehensive Serology Panel” and the “Mouse 3R Quarantine Annual SOPF PCR Panel” from IDEXX BioAnalytics. The assays were performed on pooled samples, a microorganism was considered present, if it was identified through at least one of the two independent methodologies. **The hygiene standard of conventional SPF mice was assessed and reported by the mouse vendor (Taconic Biosciences).

## References

1. T. A. E. Platts-Mills, The allergy epidemics: 1870-2010, Journal of Allergy and Clinical Immunology 136, 3–13 (2015).

2. L. Henriksen, J. Simonsen, A. Haerskjold, M. Linder, H. Kieler, S. F. Thomsen, L. G. Stensballe, Incidence rates of atopic dermatitis, asthma, and allergic rhinoconjunctivitis in Danish and Swedish children, J Allergy Clin Immunol 136, 360–366.e2 (2015).

3. C. Ober, T.-C. Yao, The genetics of asthma and allergic disease: a 21st century perspective: Genetics of asthma and allergy, Immunological Reviews 242, 10–30 (2011).

4. D. P. Strachan, Hay fever, hygiene, and household size, BMJ 299, 1259–1260 (1989).

5. D. P. Strachan, Family size, infection and atopy: the first decade of the “hygiene hypothesis,” Thorax 55 Suppl 1, S2–10 (2000).

6. C. Braun-Fahrländer, M. Gassner, L. Grize, U. Neu, F. H. Sennhauser, H. S. Varonier, J. C. Vuille, B. Wüthrich, Prevalence of hay fever and allergic sensitization in farmer’s children and their peers living in the same rural community. SCARPOL team. Swiss Study on Childhood Allergy and Respiratory Symptoms with Respect to Air Pollution, Clin Exp Allergy 29, 28–34 (1999).

7. M. Kilpeläinen, E. O. Terho, H. Helenius, M. Koskenvuo, Farm environment in childhood prevents the development of allergies, Clin Exp Allergy 30, 201–208 (2000).

8. Von Ehrenstein, Von Mutius, Illi, Baumann, Böhm, Von Kries, Reduced risk of hay fever and asthma among children of farmers: Hay fever and asthma in farmers’ children, Clinical & Experimental Allergy 30, 187–193 (2000).

9. Von Mutius, Braun-Fahrländer, Schierl, Riedler, Ehlermann, Maisch, Waser, Nowak, Exposure to endotoxin or other bacterial components might protect against the development of atopy: Endotoxin exposure protection, Clinical & Experimental Allergy 30, 1230–1234 (2000).

10. E. von Mutius, The microbial environment and its influence on asthma prevention in early life, Journal of Allergy and Clinical Immunology 137, 680–689 (2016).

11. M. Waser, K. B. Michels, C. Bieli, H. Flöistrup, G. Pershagen, E. von Mutius, M. Ege, J. Riedler, D. Schram-Bijkerk, B. Brunekreef, M. van Hage, R. Lauener, C. Braun-Fahrländer, the PARSIFAL Study team, Inverse association of farm milk consumption with asthma and allergy in rural and suburban populations across Europe, Clin Exp Allergy 37, 661–670 (2007).

12. M. J. Schuijs, M. A. Willart, K. Vergote, D. Gras, K. Deswarte, M. J. Ege, F. B. Madeira, R. Beyaert, G. van Loo, F. Bracher, E. von Mutius, P. Chanez, B. N. Lambrecht, H. Hammad, Farm dust and endotoxin protect against allergy through A20 induction in lung epithelial cells, Science 349, 1106–1110 (2015).

13. S. F. Bloomfield, R. Stanwell-Smith, R. W. R. Crevel, J. Pickup, Too clean, or not too clean: the Hygiene Hypothesis and home hygiene, Clin Exp Allergy 36, 402–425 (2006).

14. C. Brooks, N. Pearce, J. Douwes, The hygiene hypothesis in allergy and asthma: an update, *Current Opinion in Allergy* & Clinical Immunology 13, 70–77 (2013).

15. J. Douwes, Does environmental endotoxin exposure prevent asthma?, Thorax 57, 86–90 (2002).

16. A. H. Liu, J. R. Murphy, Hygiene hypothesis: Fact or fiction?, Journal of Allergy and Clinical Immunology 111, 471–478 (2003).

17. P. S. Thorne, A. Mendy, N. Metwali, P. Salo, C. Co, R. Jaramillo, K. M. Rose, D. C. Zeldin, Endotoxin Exposure: Predictors and Prevalence of Associated Asthma Outcomes in the United States, Am J Respir Crit Care Med 192, 1287–1297 (2015).

18. S. P. Rosshart, J. Herz, B. G. Vassallo, A. Hunter, M. K. Wall, J. H. Badger, J. A. McCulloch, D. G. Anastasakis, A. A. Sarshad, I. Leonardi, N. Collins, J. A. Blatter, S.-J. Han, S. Tamoutounour, S. Potapova, M. B. Foster St. Claire, W. Yuan, S. K. Sen, M. S. Dreier, B. Hild, M. Hafner, D. Wang, I. D. Iliev, Y. Belkaid, G. Trinchieri, B. Rehermann, Laboratory mice born to wild mice have natural microbiota and model human immune responses, Science 365, eaaw4361 (2019).

19. S. P. Rosshart, B. G. Vassallo, D. Angeletti, D. S. Hutchinson, A. P. Morgan, K. Takeda, H. D. Hickman, J. A. McCulloch, J. H. Badger, N. J. Ajami, G. Trinchieri, F. Pardo-Manuel de Villena, J. W. Yewdell, B. Rehermann, Wild Mouse Gut Microbiota Promotes Host Fitness and Improves Disease Resistance, Cell 171, 1015–1028.e13 (2017).

20. L. K. Beura, S. E. Hamilton, K. Bi, J. M. Schenkel, O. A. Odumade, K. A. Casey, E. A. Thompson, K. A. Fraser, P. C. Rosato, A. Filali-Mouhim, R. P. Sekaly, M. K. Jenkins, V. Vezys, W. N. Haining, S. C. Jameson, D. Masopust, Normalizing the environment recapitulates adult human immune traits in laboratory mice, Nature 532, 512–516 (2016).

21. J.-D. Lin, J. C. Devlin, F. Yeung, C. McCauley, J. M. Leung, Y.-H. Chen, A. Cronkite, C. Hansen, C. Drake-Dunn, K. V. Ruggles, K. Cadwell, A. L. Graham, P. Loke, Rewilding Nod2 and Atg16l1 Mutant Mice Uncovers Genetic and Environmental Contributions to Microbial Responses and Immune Cell Composition, Cell Host & Microbe 27, 830–840.e4 (2020).

22. F. Yeung, Y.-H. Chen, J.-D. Lin, J. M. Leung, C. McCauley, J. C. Devlin, C. Hansen, A. Cronkite, Z. Stephens, C. Drake-Dunn, Y. Fulmer, B. Shopsin, K. V. Ruggles, J. L. Round, P. Loke, A. L. Graham, K. Cadwell, Altered Immunity of Laboratory Mice in the Natural Environment Is Associated with Fungal Colonization, Cell Host & Microbe 27, 809–822.e6 (2020).

23. R. M. Green, Synergism between allergens and viruses and risk of hospital admission with asthma: case-control study, BMJ 324, 763–763 (2002).

24. C. S. Murray, Study of modifiable risk factors for asthma exacerbations: virus infection and allergen exposure increase the risk of asthma hospital admissions in children, Thorax 61, 376–382 (2006).

25. R. Sporik, S. T. Holgate, T. A. E. Platts-Mills, J. J. Cogswell, Exposure to House-Dust Mite Allergen (*Der p* I) and the Development of Asthma in Childhood: A Prospective Study, N Engl J Med 323, 502–507 (1990).

26. B. N. Lambrecht, H. Hammad, The immunology of asthma, Nat Immunol 16, 45–56 (2015).

27. C. A. Tibbitt, J. M. Stark, L. Martens, J. Ma, J. E. Mold, K. Deswarte, G. Oliynyk, X. Feng, B. N. Lambrecht, P. De Bleser, S. Nylén, H. Hammad, M. Arsenian Henriksson, Y. Saeys, J. M. Coquet, Single-Cell RNA Sequencing of the T Helper Cell Response to House Dust Mites Defines a Distinct Gene Expression Signature in Airway Th2 Cells, Immunity 51, 169–184.e5 (2019).

28. B. D. Hondowicz, D. An, J. M. Schenkel, K. S. Kim, H. R. Steach, A. T. Krishnamurty, G. J. Keitany, E. N. Garza, K. A. Fraser, J. J. Moon, W. A. Altemeier, D. Masopust, M. Pepper, Interleukin-2-Dependent Allergen-Specific Tissue-Resident Memory Cells Drive Asthma, Immunity 44, 155–166 (2016).

29. D. Artis, H. Spits, The biology of innate lymphoid cells, Nature 517, 293–301 (2015).

30. L. Guo, Y. Huang, X. Chen, J. Hu-Li, J. F. Urban, W. E. Paul, Innate immunological function of TH2 cells in vivo, Nat Immunol 16, 1051–1059 (2015).

31. R. M. Maizels, Regulation of immunity and allergy by helminth parasites, Allergy 75, 524–534 (2020).

32. G. A. W. Rook, V. Adams, R. Palmer, L. R. Brunet, J. Hunt, R. Martinelli, Mycobacteria and other environmental organisms as immunomodulators for immunoregulatory disorders, Springer Seminars in Immunopathology 25, 237–255 (2004).

33. Y. M. Sjögren, M. C. Jenmalm, M. F. Böttcher, B. Björkstén, E. Sverremark-Ekström, Altered early infant gut microbiota in children developing allergy up to 5 years of age, Clinical & Experimental Allergy 39, 518–526 (2009).

34. M. J. Ege, M. Mayer, A.-C. Normand, J. Genuneit, W. O. C. M. Cookson, C. Braun-Fahrländer, D. Heederik, R. Piarroux, E. von Mutius, Exposure to Environmental Microorganisms and Childhood Asthma, N Engl J Med 364, 701–709 (2011).

35. T. A. E. Platts-Mills, E. Erwin, P. Heymann, J. Woodfolk, Is the hygiene hypothesis still a viable explanation for the increased prevalence of asthma?, Allergy 60, 25–31 (2005).

36. S.-N. Yang, C.-C. Hsieh, H.-F. Kuo, M.-S. Lee, M.-Y. Huang, C.-H. Kuo, C.-H. Hung, The Effects of Environmental Toxins on Allergic Inflammation, Allergy Asthma Immunol Res 6, 478 (2014).

## References

37. T. R. Abrahamsson, H. E. Jakobsson, A. F. Andersson, B. Björkstén, L. Engstrand, M. C. Jenmalm, Low diversity of the gut microbiota in infants with atopic eczema, J Allergy Clin Immunol 129, 434–440, 440.e1-2 (2012).

38. M. Wang, C. Karlsson, C. Olsson, I. Adlerberth, A. E. Wold, D. P. Strachan, P. M. Martricardi, N. Åberg, M. R. Perkin, S. Tripodi, A. R. Coates, B. Hesselmar, R. Saalman, G. Molin, S. Ahrné, Reduced diversity in the early fecal microbiota of infants with atopic eczema, Journal of Allergy and Clinical Immunology 121, 129–134 (2008).

39. B. M. Henrick, L. Rodriguez, T. Lakshmikanth, C. Pou, E. Henckel, A. Olin, J. Wang, J. Mikes, Z. Tan, Y. Chen, A. M. Ehrlich, A. K. Bernhardsson, C. H. Mugabo, Y. Ambrosiani, A. Gustafsson, S. Chew, H. K. Brown, J. Prambs, K. Bohlin, R. D. Mitchell, M. A. Underwood, J. T. Smilowitz, J. B. German, S. A. Frese, P. Brodin, Bifidobacteria-mediated immune system imprinting early in life (Immunology, 2020; http://biorxiv.org/lookup/doi/10.1101/2020.10.24.353250).

40. B. Björkstén, E. Sepp, K. Julge, T. Voor, M. Mikelsaar, Allergy development and the intestinal microflora during the first year of life, Journal of Allergy and Clinical Immunology 108, 516–520 (2001).

41. M. Kalliomäki, P. Kirjavainen, E. Eerola, P. Kero, S. Salminen, E. Isolauri, Distinct patterns of neonatal gut microflora in infants in whom atopy was and was not developing, Journal of Allergy and Clinical Immunology 107, 129–134 (2001).

42. N. Blümer, S. Sel, S. Virna, C. C. Patrascan, S. Zimmermann, U. Herz, H. Renz, H. Garn, Perinatal maternal application of Lactobacillus rhamnosus GG suppresses allergic airway inflammation in mouse offspring, Clin Exp Allergy 37, 348–357 (2007).

43. J. Debarry, H. Garn, A. Hanuszkiewicz, N. Dickgreber, N. Blümer, E. von Mutius, A. Bufe, S. Gatermann, H. Renz, O. Holst, H. Heine, Acinetobacter lwoffii and Lactococcus lactis strains isolated from farm cowsheds possess strong allergy-protective properties, Journal of Allergy and Clinical Immunology 119, 1514–1521 (2007).

44. K. Karimi, M. D. Inman, J. Bienenstock, P. Forsythe, *Lactobacillus reuteri* -induced Regulatory T cells Protect against an Allergic Airway Response in Mice, Am J Respir Crit Care Med 179, 186–193 (2009).

45. A. Trompette, E. S. Gollwitzer, K. Yadava, A. K. Sichelstiel, N. Sprenger, C. Ngom-Bru, C. Blanchard, T. Junt, L. P. Nicod, N. L. Harris, B. J. Marsland, Gut microbiota metabolism of dietary fiber influences allergic airway disease and hematopoiesis, Nat Med 20, 159–166 (2014).

46. H. J. McSorley, M. T. O’Gorman, N. Blair, T. E. Sutherland, K. J. Filbey, R. M. Maizels, Suppression of type 2 immunity and allergic airway inflammation by secreted products of the helminth Heligmosomoides polygyrus, Eur J Immunol 42, 2667–2682 (2012).

47. C. J. C. Johnston, D. J. Smyth, R. B. Kodali, M. P. J. White, Y. Harcus, K. J. Filbey, J. P. Hewitson, C. S. Hinck, A. Ivens, A. M. Kemter, A. O. Kildemoes, T. Le Bihan, D. C. Soares, S. M. Anderton, T. Brenn, S. J. Wigmore, H. V. Woodcock, R. C. Chambers, A. P. Hinck, H. J. McSorley, R. M. Maizels, A structurally distinct TGF-β mimic from an intestinal helminth parasite potently induces regulatory T cells, Nat Commun 8, 1741 (2017).

48. C. Haluszczak, A. D. Akue, S. E. Hamilton, L. D. S. Johnson, L. Pujanauski, L. Teodorovic, S. C. Jameson, R. M. Kedl, The antigen-specific CD8+ T cell repertoire in unimmunized mice includes memory phenotype cells bearing markers of homeostatic expansion, Journal of Experimental Medicine 206, 435–448 (2009).

49. R. W. Nelson, D. Beisang, N. J. Tubo, T. Dileepan, D. L. Wiesner, K. Nielsen, M. Wüthrich, B. S. Klein, D. I. Kotov, J. A. Spanier, B. T. Fife, J. J. Moon, M. K. Jenkins, T Cell Receptor Cross-Reactivity between Similar Foreign and Self Peptides Influences Naive Cell Population Size and Autoimmunity, Immunity 42, 95–107 (2015).

50. C.-C. Chen, T. Kobayashi, K. Iijima, F.-C. Hsu, H. Kita, IL-33 dysregulates regulatory T cells and impairs established immunologic tolerance in the lungs, Journal of Allergy and Clinical Immunology 140, 1351–1363.e7 (2017).

51. J. M. Han, D. Wu, H. C. Denroche, Y. Yao, C. B. Verchere, M. K. Levings, IL-33 Reverses an Obesity-Induced Deficit in Visceral Adipose Tissue ST2 ^+^ T Regulatory Cells and Ameliorates Adipose Tissue Inflammation and Insulin Resistance, J.I. 194, 4777–4783 (2015).

52. W. Kuswanto, D. Burzyn, M. Panduro, K. K. Wang, Y. C. Jang, A. J. Wagers, C. Benoist, D. Mathis, Poor Repair of Skeletal Muscle in Aging Mice Reflects a Defect in Local, Interleukin-33-Dependent Accumulation of Regulatory T Cells, Immunity 44, 355–367 (2016).

53. A. Vasanthakumar, K. Moro, A. Xin, Y. Liao, R. Gloury, S. Kawamoto, S. Fagarasan, L. A. Mielke, S. Afshar-Sterle, S. L. Masters, S. Nakae, H. Saito, J. M. Wentworth, P. Li, W. Liao, W. J. Leonard, G. K. Smyth, W. Shi, S. L. Nutt, S. Koyasu, A. Kallies, The transcriptional regulators IRF4, BATF and IL-33 orchestrate development and maintenance of adipose tissue-resident regulatory T cells, Nat Immunol 16, 276–285 (2015).

